# Investigating the impact of obesity as an immunological instigator on young-onset arthritis using perturbation-based simulation

**DOI:** 10.64898/2026.01.29.702692

**Authors:** Tyler Hatstadt, Michael Bryan

## Abstract

**Purpose:** Young-onset inflammatory arthritis (YOA; disease onset at age <25) incidence has risen since 1990, coinciding with the childhood obesity epidemic. Traditional epidemiology cannot easily quantify obesity’s contribution because early-life exposures can precede clinical onset by years and may be poorly measured at diagnosis. We developed a perturbation-based simulation to estimate young-age inflammatory arthritis burden by calibrating to age-stratified RA estimates from GBD (GBD does not report JIA) under varying obesity scenarios - an approach that allows counterfactual testing, difficult to achieve in observational studies.

**Methods:** We built a Monte Carlo model generating arthritis prevalence estimates for 1,000,000 individuals. The model incorporated published odds ratios: BMI per SD, smoking, HLA-DR shared epitope allele dose (0/1/2), and interactions. We systematically perturbed average BMI in the <25 stratum (weighted towards adolescents and young adults) from 25-29 kg/m^2^ as a stress-test while holding other factors constant, then compared predicted prevalence against Global Burden of Disease data. Each scenario ran 2,500 iterations to propagate parameter uncertainty.

**Results:** Our model predicted the current YOA prevalence of 0.07% (observed: 0.06%, 95% CI: 0.05%-0.07%). Under perturbation analysis, each unit increase in average BMI yielded an additional 0.005% (95% CI: 0.0025%-0.0075%) in YOA cases - small but meaningful given the rarity of young-onset inflammatory arthritis. The relationship was locally linear. Significantly, the model saw that returning average BMI toward 1990 levels (<26) predicted around a 30% drop in BMI-attributable diagnoses and a 3% decrease in YOA prevalence.

**Conclusions:** Perturbation modeling identifies childhood obesity as a potentially modifiable driver of young-onset RA, accounting for upwards of 5% of prevalence increases since 1990. This approach uniquely enables testing of prevention scenarios: our model predicts that lowering average BMI by one to two units over the next decade could prevent 3-5% of YOA cases in <25 under the modeled scenarios. These estimates provide a quantitative basis for incorporating arthritis prevention in childhood and adolescent obesity intervention cost-effectiveness analyses.

## Introduction

The reported incidence of autoimmune young-onset inflammatory arthritis (YOA) in adolescents and young adults has steadily increased since 1990 (e.g., increase from ∼4.98 to 5.41 per 100,000 in adolescent/young-adult rheumatoid arthritis incidence from 1990 to 2019 in GBD-based estimates). These conditions, specifically rheumatoid arthritis (RA), juvenile idiopathic arthritis (JIA; referred to generally as JA), and their various subtypes, are temporally coincident with rising early-life and adolescent obesity rates. While obesity promotes systemic inflammation, its specific contribution to autoimmune arthritis development remains unknown. Most observational studies assess BMI in adulthood^1 2^, often near the time of diagnosis, limiting their ability to evaluate effects mediated by early-life exposures with long lag times. We used simulation modeling to estimate the plausible contribution of childhood obesity to young-onset inflammatory arthritis prevalence.

Dietary and environmental changes in the U.S. over the past 30 years are driving the significant increase in rates of obesity and average BMI, especially in younger people. Approximately 40% of U.S. adults meet the criteria for obesity.^3^ Available GBD estimates show smooth monotonic increases over time, although these modeled series may obscure true discontinuities arising from changes in diagnostic practices.^15^

Obesity is traditionally viewed as an epigenetic/lifestyle-dependent condition. In contrast, RA and several JA subtypes are strongly influenced by genetic susceptibility. While research on JA has only recently begun, certain subtypes of JA and RA share similarities in both symptoms and onset. Many HLA-DR4 haplotypes carry HLA-DRB1 ‘shared epitope’ (SE) alleles, which are among the strongest common genetic risk factors for RA. The interactions of these genes and immune activation can and often are regulated through the epigenome, thus allowing specific environmental factors to increase the likelihood of RA development. Some of these non-genetic conditions, such as smoking, are well-known risk factors. Smoking is often associated with the more severe side of symptoms in those afflicted with seropositive RA. Obesity has also been linked to a higher likelihood of developing rheumatoid arthritis in recent studies.

A widely supported model of disease onset in RA posits individual sites of inflammation, typically initiated on mucosal surfaces, where the immune reaction is systemically propagated over time, resulting in chronic inflammation in the joints. This method is greatly influenced by factors such as smoking, which damages mucosal surfaces in the lungs. However, there are several distinct mechanisms involving both the innate and adaptive systems that are implicated in RA pathogenesis, notably seronegative RA. An emerging model driven by adipose tissue appears to be increasingly relevant in younger populations, particularly in women. This process represents one plausible pathway among several, and their relative contribution likely varies across disease subtypes and individuals.^4^

Over the past few years, adipose has been repeatedly recognized as an active endocrine and immune organ. Adipose tissue secretes cytokines (adipokines) like TNF-alpha, IL-6, IL-1beta, leptin, resistin, and free fatty acids (FFAs). In individuals with an abnormally high BMI, these cytokines cause the near constant activation of NF-kappaB and JAK/STAT, a systemic M1 macrophage polarization, and the suppression of regulatory T cells (Tregs). This phenomenon supports the claim that high adiposity can lead to a body-wide condition of chronic low-grade innate inflammation without infection.^5^

In parallel, monocytes and macrophages undergo epigenetic reprogramming, changing histone acetylation, DNA methylation, and chromatin accessibility at cytokine genes. These alterations then create a semblance of trained immunity, where future inflammatory responses become exaggerated and self-sustaining. A small trigger produces a massive cytokine output, and resolution mechanisms become inherently inefficient, gradually approaching a state of locked-in hyper-reactive innate immunity. This can occur in the absence of antigen-specific adaptive activation.^6^

Because the joint synovium is highly vascular and cytokine-sensitive, the circulating adipokines trigger signals in macrophages and fibroblast-like synoviocytes (FLS), which induce several reactions that create joint inflammation before adaptive autoimmunity is established. The FLS become hyperproliferative once exposed to adipokines, lose contact inhibition, and become invasive. They begin secreting MMPs, RANKL, and certain other cytokines, creating a pannus-like aggressive synovium. The macrophages and FLS also begin osteoclastogenesis without needing ACPA/anti-CCP, possibly explaining how seronegative RA can still be erosive.^7^

Only after chronic innate synovitis is established does the adaptive immune system fully activate, and low levels of RF and ACPA/anti-CCP will be produced - they are downstream, not initiating. The pathway is reversed from classic RA, though not exclusive: first beginning with the innate system, then synovitis occurs, leading to tissue damage that triggers adaptive autoimmunity. Taken together, these interactions support a model in which obesity-driven innate immune dysregulation can initiate and sustain synovial inflammation independently of adaptive autoimmunity.^8^

While JA is a genetically unique subtype of autoimmune arthritis compared to RA, the mechanism described above has the potential to be activated regardless of genotype or phenotype. Importantly, some subtypes of JA are autoinflammatory rather than autoimmune, plausibly supporting this distinction. Due to the obesity pathway’s activation of the innate immune system, the body can produce symptoms indicative of either RA or JA without producing detectable autoantibodies. The shared outcome is chronic joint inflammation, although underlying immunopathology may differ. In patients with the appropriate genetic and epigenetic status, the adaptive immune system may be triggered after symptoms have already begun to show, potentially explaining the presence of certain seronegative RA and JA cases.

Understanding obesity’s specific contribution to autoimmune arthritis onset is difficult in typical observational studies, even after adjusting for confounding variables. Many cohorts are too small or poorly phenotyped to see any statistically significant impact. To offset this, we built a Monte Carlo simulation-based perturbation model of BMI distributions and ran several counterfactual scenarios difficult to obtain in real datasets. On 1,000,000 simulated patients, each risk factor and its contribution towards disease outcome per patient was tracked, including smoking status, genotype, BMI, age, and interaction terms. The process allowed for the approximate estimation of BMI’s population-level impact on the development of RA/JA with respect to age. Because the model is designed to estimate relative population-level shifts under counterfactual BMI scenarios, results should be interpreted as plausible contributions under assumed causal effects rather than direct causal estimates.

## Methods & Materials

### Dataset

The Institute of Health Metrics and Evaluation (IHME) provides a publicly accessible tool known as the GBD Study 2023.^15^ Covering 204 countries and territories over 30 years, the GBD Study 2023 integrates data from tens of thousands of epidemiological sources, including national health surveys, disease registries, hospital records, and vital registration systems, collectively representing billions of individual observations worldwide. The GBD 2023 is one of the most comprehensive and widely used sources of global disease burden estimates, including both RA and obesity. The GBD combines heterogeneous data sources using Bayesian meta-regression to generate internally weighted estimates. The GBD Study 2023 tool generates 95% uncertainty intervals for each data point determined from 1,000 draws, dependent on internal weights on multiple data sources. Every value has an associated uncertainty interval with bounds defined by the 25th and 975th order statistics, corresponding to the 2.5th and 97.5th percentiles.

Since the GBD does not include statistics on JA, for simulation purposes, population-level prevalence of RA per 100,000 was used as a steady-state approximation of population-level disease probability under stable incidence and survival assumptions. To characterize long-term trends, simple regression models were fit to RA incidence over time, and model fit was determined using standard regression diagnostics, including the coefficient of determination (R^2^) and residuals. The linear model demonstrated the highest explanatory power with little to no systematic residual patterns. Residuals showed no substantial autocorrelation. The linear model was thus selected as the best-fitting model.

### Inputs for Simulation

Odds ratios (ORs) were derived from meta-analyses of carefully selected, peer-reviewed external cohort studies for the effects of BMI per standard deviation (SD), smoking status, genotype, and age on arthritis likelihood. To avoid over-interpretation of any single study, parameter values were chosen to reflect central tendencies within published ranges rather than exact estimates from individual cohorts. While ORs are usually derived from incidence studies, the model operates under a rare-disease approximation, which is plausible given the low population prevalence of RA and JA. A forest plot was drawn to visually show the statistical significance of each risk factor, where odds ratios whose 95% confidence intervals (CIs) crossed unity were noted as not statistically significant. These odds ratios were then converted to logistic regression coefficients and used to parameterize a probabilistic risk model. Additionally, a single baseline prevalence (GBD 2023, all ages, both sexes, U.S.-specific) was used to set the intercept; GBD age-stratified prevalence estimates were used only for calibration comparisons. Baseline prevalence and demographic distributions were U.S.-specific; effect sizes were drawn from published multi-cohort meta-analyses.

### Simulation Approach

Using these coefficients, 1,000,000 patients were simulated with Bernoulli trials, mapping linear predictors to probabilities with a standard logistic sigmoid function to include both continuous and binary values (e.g., BMI versus smoking status). Each patient’s OR was generated according to a Gaussian distribution in log space, representing the upper and lower bounds of the respective confidence intervals.

Standard errors were inferred from reported 95% confidence intervals assuming approximate normality in log-odds space. A single baseline prevalence was used to isolate relative risk shifts driven by covariates, with age effects modeled through stratification and interaction terms. Assuming baseline arthritis risk based on the GBD’s RA prevalence for all ages, a risk value was calculated per patient, and disease outcome was generated using the Monte Carlo method. Patients were then stratified by age group (15-24, 25-55, 55+). Each stratum’s predicted RA prevalence estimates were calibrated and checked for internal consistency against GBD age-stratified prevalence estimates. This approach allowed both qualitative and quantitative comparison of disease risk across stratified subpopulations.

## Simulation Calibration

To explore age-dependent scaling, the function for determining the OR for BMI per SD used an exponential decay function. The conditions of the function were that the maximum value of the OR confidence interval (1.26) decayed to the minimum value (1.01) with age, and the average value of the function remained the point estimate (1.13). A sensitivity analysis was performed on the decay constant k to show significant or insignificant changes in obesity prevalence among patients with arthritis. An exponential form was chosen to represent age-dependent scaling while minimizing model complexity. This ensured a stronger impact in younger patients and a lesser but still notable impact in middle-aged and older patients while remaining consistent with sampled confidence intervals.

### Simulation Uncertainty

The Monte Carlo method allowed for counterfactual testing and perturbation analysis. The center for the Gaussian distribution of BMI was systematically perturbed as a stress-test in the youngest age group from 25 to 29 units, normalized around the assumed U.S. average of 27 units. The <25 stratum is weighted toward adolescents and young adults based on population age structure, which was distributed according to U.S. age demographics; BMI was modeled on the adult BMI scale (kg/m^2^) as a stylized proxy for adiposity distribution in this stratum. Using a bootstrap resampling method on 1,000,000 simulated patients over 2,500 trials, confidence intervals were drawn for each age group for arthritis risk, arthritis prevalence, average BMI in patients with arthritis, and obesity prevalence in patients with arthritis.

## Results

**Figure 1.**
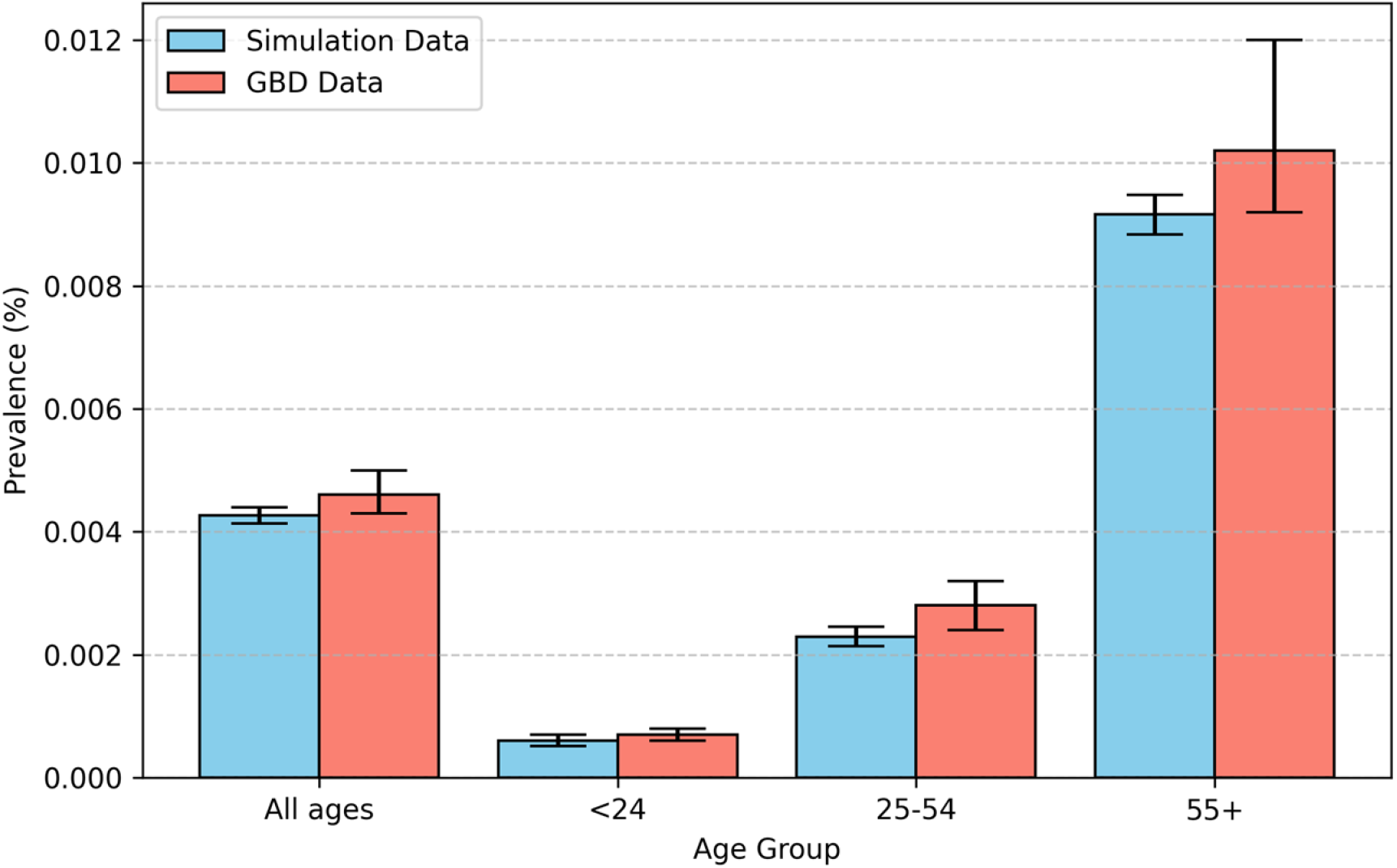
Simulated outputs were assessed for calibration consistency with observed RA prevalence (used as a proxy for young-onset inflammatory arthritis) across age strata in the GBD dataset^15^. Simulated prevalence estimates closely matched GBD age-stratified prevalence, with overlapping 95% uncertainty intervals across all age groups. Median simulated prevalence values fell within the corresponding GBD uncertainty bounds, with no systematic deviation by age. This agreement supported the use of the model for subsequent risk factor and counterfactual analyses.

**Figure 2.**
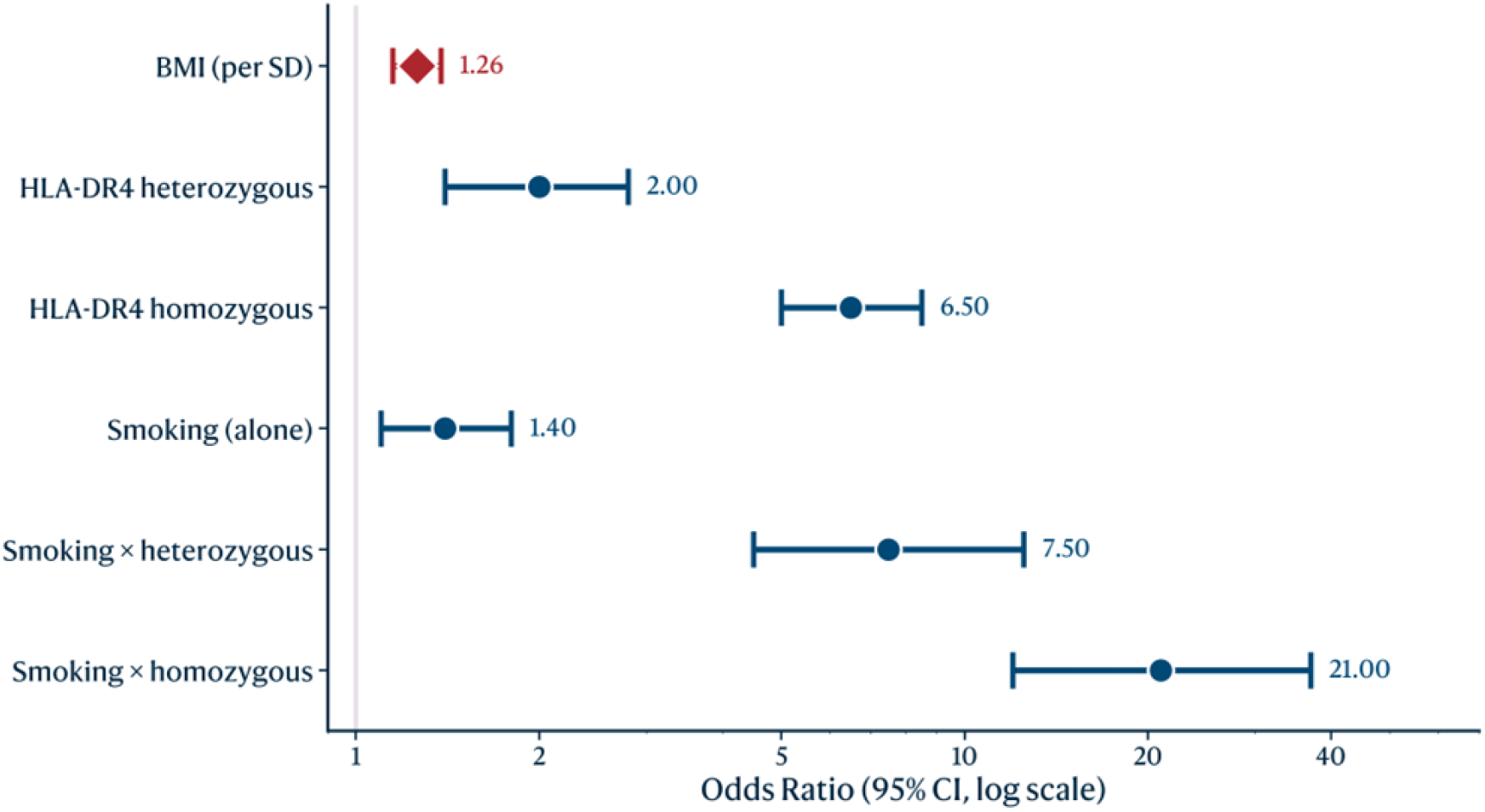
Model parameterization incorporated literature-derived odds ratios for BMI per standard deviation, HLA-DR shared epitope allele dose, smoking status as a binary feature, and smoking-genotype interaction terms. Genetic and smoking-related factors exhibited larger per-exposure effect sizes than BMI. Alcohol consumption showed no statistically significant association with autoimmune arthritis risk (OR = 0.8-1.1) and was therefore excluded from subsequent modeling. These parameters were used to construct the probabilistic risk model. ^9 10 11 12^

**Figure 3.**
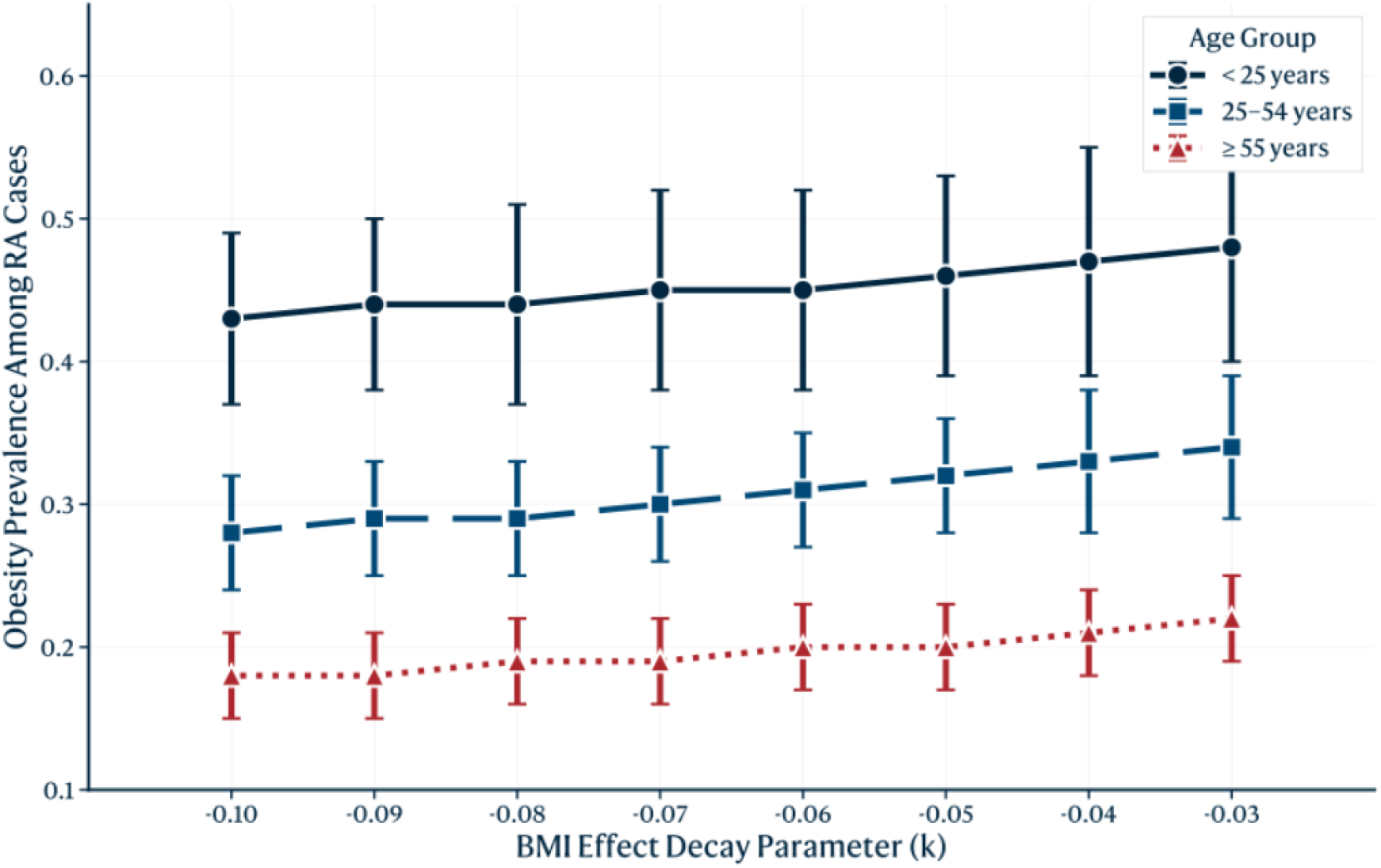
Sensitivity analyses were performed to evaluate model dependence on assumptions regarding the age-dependent scaling of BMI effects. Varying the exponential decay constant across the tested range produced negligible changes in estimated obesity prevalence among individuals with arthritis. Model outputs were stable across the tested parameter space.

**Figure 4.**
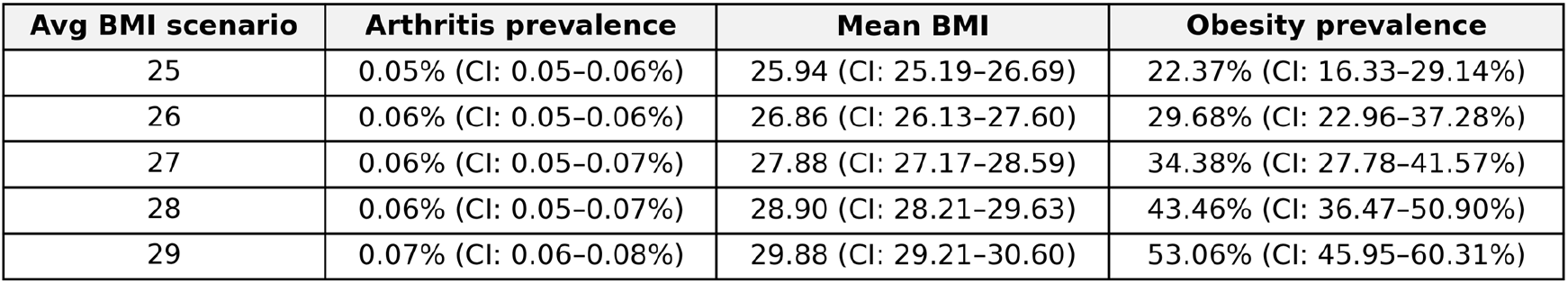
Counterfactual perturbation analyses were conducted by holding all risk factors constant and systematically varying mean BMI in the <25 stratum. Changes in mean BMI were associated with proportional changes in modeled arthritis prevalence, approximating a locally linear relationship over the tested BMI range. Bootstrap resampling produced narrow confidence intervals across trials and age groups. These results should be interpreted as estimates of plausible contribution rather than causal effect, given the model’s reliance on literature-derived parameters and population-level assumptions.

Together, these results establish internal consistency, parameter stability, and sensitivity to population-level BMI perturbations, motivating interpretation of modeled BMI effects as a plausible contributor to young-onset disease risk. Simulation does not establish causality, but can provide a quantitative framework for counterfactual analysis and hypothesis generation.

## Discussion

This study used a simulation-based framework to evaluate the plausible contribution of childhood and adolescent obesity to the risk of young-onset inflammatory arthritis. Across multiple calibration and sensitivity analyses, the model consistently reproduced age-stratified prevalence. Results demonstrated that population-level variation in BMI was associated with proportional changes in modeled disease risk, particularly in younger age groups. Although genetic susceptibility and smoking exhibited larger effect sizes, BMI remained a stable, non-negligible contributor under a range of assumptions. Notably, smoking is an individually modifiable exposure with a large effect size, whereas genotype frequencies are relatively stable at the population level; in contrast, BMI is both population-shifting and difficult to modify sustainably, which may amplify its public health relevance despite smaller per-person effect estimates. Together, these findings support a role for early-life adiposity as a modifying factor in autoimmune arthritis risk, helping to reconcile strong mechanistic evidence with the modest and inconsistent associations observed in traditional epidemiological studies.

The age-dependent scaling applied to BMI effects in the simulation was motivated by known biological constraints on obesity-driven immune dysregulation. Excess adiposity promotes innate immune activation through adipokine signaling, macrophage polarization, and trained immunity, processes that are particularly influential during periods of immune development and maturation. Innate immune plasticity is highest during early childhood and adolescence and therefore remains highly susceptible to epigenetic imprinting. Beyond early adulthood, typically by the mid-20s, hematopoietic stem cell output slows and innate immune cells exhibit reduced epigenetic responsiveness. As age increases, additional drivers of arthritis risk, such as cumulative joint stress, immunosenescence, and stochastic immune activation, become increasingly dominant, reducing the relative contribution of adiposity without eliminating it. Direct human evidence for the timing and magnitude of these transitions remains limited, and the functional form should be interpreted as a biologically motivated approximation rather than a measured trajectory. Modeling this transition as a smooth exponential decay captures a biologically plausible shift from early-life susceptibility to multifactorial adult disease, while remaining consistent with reported confidence intervals for BMI-associated risk. The stability of model outputs across a wide range of decay constants further suggests that the results are not artifacts of functional form, but reflect a robust relationship between early adiposity and disease risk.^13 14^

Traditional epidemiological approaches are poorly suited to estimating the contribution of early-life adiposity to autoimmune arthritis across the life course. Most large observational cohorts measure BMI at or near the time of diagnosis, implicitly assuming contemporaneous exposure-outcome relationships, while being unable to capture critical exposure windows occurring potentially decades earlier. This temporal misalignment introduces substantial attenuation bias, particularly when the exposure exerts its effects through immune programming rather than immediate pathophysiology. Additionally, BMI is a dynamic, population-shifting variable with relatively small individual-level effect sizes, making its contribution difficult to detect against stronger, static, and more readily measurable risk factors such as genotype or smoking. Residual confounding, exposure misclassification, and survivorship bias further obscure modest multiplicative effects, especially when risk is distributed across heterogeneous subpopulations rather than concentrated in narrowly defined high-risk groups.

To our knowledge, early-life adiposity has not been systematically evaluated in large longitudinal cohorts with sufficient follow-up to assess its contribution to late-onset rheumatoid arthritis pathogenesis. Most available cohort studies measure BMI contemporaneously with diagnosis or in mid-to late adulthood, limiting their ability to detect effects mediated through early immune programming. Given the capacity of the innate immune system for durable epigenetic imprinting during development, early-life adiposity may plausibly influence inflammatory set points that persist into later adulthood, thereby contributing to arthritis risk even when clinical onset occurs decades later.

The simulation framework employed addresses these limitations by decoupling biological plausibility from observational tractability. By explicitly modeling BMI as a population-level perturbation rather than a fixed individual exposure, the model captures risk shifts that emerge only when small effects act coherently across large cohorts. Importantly, the model does not rely on direct causal inference from BMI to disease, but instead integrates empirically derived effect sizes within a probabilistic structure that preserves uncertainty and interaction effects. In doing so, the simulation provides a mechanistically informed explanation for why obesity may materially contribute to young-onset disease risk despite appearing weak or inconsistent in conventional epidemiological analyses, which frequently emphasize adult exposures and outcomes.

There are several important limitations worth noting. First, the simulation framework relies on literature-derived odds ratios and population-level assumptions rather than individual longitudinal data, and therefore cannot independently establish causality between adiposity and autoimmune arthritis onset. While uncertainty was explicitly incorporated through probabilistic sampling and sensitivity analyses, unmeasured confounders such as diet composition, physical activity, socioeconomic status, and early-life infections were not directly modeled and may modify observed relationships. Second, juvenile arthritis and rheumatoid arthritis were jointly considered under a unified modeling framework despite their biological heterogeneity; although this was motivated by shared inflammatory mechanisms and the absence of comprehensive JA prevalence data, disease-specific differences may not be fully captured. Third, the use of a rare-disease approximation and a single baseline prevalence simplifies complex incidence, remission, and survival dynamics that may vary across populations and time. Finally, the age-dependent scaling of BMI effects, while biologically motivated and robust to parameter variation, represents a stylized approximation of immune maturation rather than a direct measurement of immune programming. As such, results should be interpreted as estimates of plausible population-level contribution rather than precise effect sizes or individual risk predictions.

Future work could extend this framework by integrating longitudinal cohort data with early-life BMI trajectories, enabling direct testing of the timing and persistence of obesity-associated immune programming. Introducing an explicit lag structure would allow modeling of delayed incidence and prevalence shifts across age groups. Incorporating additional environmental exposures, such as diet composition, physical activity, infection history, and socioeconomic factors, would allow more refined modeling of interacting risk pathways. From a translational perspective, these findings suggest that population-level interventions targeting childhood obesity may have downstream effects on autoimmune disease burden beyond metabolic outcomes alone, if the modeled associations reflect causal pathways. While the present results do not imply determinism at the individual level, they motivate consideration of early-life adiposity as a modifiable component of autoimmune arthritis risk, warranting further investigation in prospective and mechanistic studies.

## Notes

### Competing Interest Statement

The authors have declared no competing interest.

## Bibliography

1. Wesley A, Bengtsson C, Elkan AC, Klareskog L, Alfredsson L, Wedrén S, et al. Association between body mass index and anti-citrullinated protein antibody-positive and anti-citrullinated protein antibody-negative rheumatoid arthritis: results from a population-based case–control study. Arthritis Care Res (Hoboken). 2013;65(1):107–112. doi:10.1002/acr.21749.

2. Qin B, Yang M, Fu H, Ma N, Wei T, Tang Q, Hu Z, Liang Y, Yang Z, Zhong R. Body mass index and the risk of rheumatoid arthritis: a systematic review and dose-response meta-analysis. Arthritis Res Ther. 2015;17:86. doi:10.1186/s13075-015-0601-x.

3. Centers for Disease Control and Prevention (CDC). Prevalence of obesity and severe obesity among adults: United States, 2021–2023. NCHS Data Brief. 2024;(508).

4. Holers VM, Demoruelle MK, Kuhn KA, et al. Rheumatoid arthritis and the mucosal origins hypothesis: protection turns to destruction. Nat Rev Rheumatol. 2018;14(9):542–557. doi:10.1038/s41584-018-0070-0.

5. Winer DA, Winer S, Shen L, et al. B cells promote insulin resistance through modulation of T cells and production of pathogenic IgG antibodies. Nat Med. 2011;17(5):610–617. doi:10.1038/nm.2353.

6. Netea MG, Joosten LAB, Latz E, et al. Trained immunity: a program of innate immune memory in health and disease. Science. 2016;352(6284):aaf1098. doi:10.1126/science.aaf1098.

7. Bartok B, Firestein GS. Fibroblast-like synoviocytes: key effector cells in rheumatoid arthritis. Immunol Rev. 2010;233(1):233–255. doi:10.1111/j.0105-2896.2009.00859.x.

8. Firestein GS, McInnes IB. Immunopathogenesis of rheumatoid arthritis. Immunity. 2017;46(2):183–196. doi:10.1016/j.immuni.2017.02.006.

9. Padyukov L, Silva C, Stolt P, Alfredsson L, Klareskog L. A gene–environment interaction between smoking and shared epitope genes in HLA-DR provides a high risk of seropositive rheumatoid arthritis. Arthritis Rheum. 2004;50(10):3085–3092. doi:10.1002/art.20553.

10. Di Giuseppe D, Discacciati A, Orsini N, Wolk A. Cigarette smoking and risk of rheumatoid arthritis: a dose-response meta-analysis. Arthritis Res Ther. 2014;16(2):R61. doi:10.1186/ar4498.

11. Raychaudhuri S. Recent advances in the genetics of rheumatoid arthritis. Curr Opin Rheumatol. 2010;22(2):109–118. doi:10.1097/BOR.0b013e328336474d.

12. Jin Z, Liang L, Cui X, et al. Alcohol consumption as a preventive factor for developing rheumatoid arthritis: a dose-response meta-analysis of prospective studies. Ann Rheum Dis. 2014;73(10):1962–1967. doi:10.1136/annrheumdis-2013-204502.

13. Panda A, Arjona A, Sapey E, et al. Human innate immunosenescence: causes and consequences for immunity in old age. Trends Immunol. 2009;30(7):325–333. doi:10.1016/j.it.2009.05.004.

14. Simon AK, Hollander GA, McMichael A. Evolution of the immune system in humans from infancy to old age. Proc Biol Sci. 2015;282(1821):20143085. doi:10.1098/rspb.2014.3085.

15. Global Burden of Disease Collaborative Network. Global Burden of Disease Results (GBD Results). Seattle, WA: Institute for Health Metrics and Evaluation (IHME), University of Washington; 2024. Available from https://vizhub.healthdata.org/gbd-results/

